# Neuroaffective profiles are associated with e-cigarette use

**DOI:** 10.1101/2022.02.04.479183

**Authors:** Francesco Versace, George Kypriotakis

## Abstract

**Introduction:** Identifying the psychophysiological underpinnings of cue-induced compulsive nicotine use will provide new targets for relapse prevention treatments. We tested whether neuroaffective responses to motivationally relevant stimuli are associated with cue-induced nicotine self-administration. We hypothesized that smokers with stronger neuroaffective responses to nicotine-related cues than to pleasant stimuli (C>P) are more vulnerable to cue-induced nicotine self-administration than smokers with stronger neuroaffective responses to pleasant stimuli than to nicotine-related cues (P>C).

**Methods:** Smokers (N=36) looked at pleasant, unpleasant, neutral, and nicotine-related images signaling that an electronic nicotine delivery system (ENDS) was immediately available for use. We measured event-related potentials (a direct measure of cortical activity) and computed the amplitude of the late positive potential, a robust index of motivational salience. We used *k*-means cluster analysis to identify individuals characterized by the C>P or the P>C neuroaffective profile. We compared the ENDS use frequency in the two groups using quantile regression for counts.

**Results:** Cluster analysis assigned 18 smokers to the C>P profile and 18 smokers to the P>C profile. Smokers with the C>P neuroaffective profile used the ENDS significantly more often than smokers with the P>C profile. Significant differences in the number of puffs persisted across different quantiles.

**Conclusions:** These results support the hypothesis that individual differences in the tendency to attribute motivational salience to drug-related cues underlie vulnerability to cue-induced drug self-administration.

**Implications:** By linking brain reactivity profiles to nicotine self-administration, we identified a neuroaffective biomarker that could guide the development of personalized treatments to prevent smoking relapse.

## Introduction

The presence of cigarette-related cues (i.e., stimuli associated with smoking) can trigger smoking^1^ and relapse.^2^ Identifying the psychophysiological mechanisms that increase vulnerability to cue-induced compulsive nicotine seeking will foster the development of tailored relapse-prevention interventions.

According to the incentive sensitization theory of addiction,^3^ vulnerability to cue-induced compulsive drug seeking stems from attributing high incentive salience to drug-related cues. Incentive salience refers to the motivational properties that make a stimulus wanted.^4^ Repeated drug use can sensitize the mesocorticolimbic dopamine systems, biasing them to attribute high incentive salience to drug-related cues. Once cues acquire incentive salience, they capture attention, activate affective states, and motivate drug seeking. Preclinical findings showed that some animals are more vulnerable than others to sensitization, and those that attribute greater incentive salience to reward-related discrete cues are prone to cue-induced compulsive drug seeking.^5^

We proposed that individual differences in the tendency to attribute incentive salience to reward-related cues also underlie vulnerability to cue-induced drug seeking in humans.^6^ Measuring brain responses to a wide array of emotional stimuli, we previously showed that individuals with larger neuroaffective responses to drug-related cues than to pleasant stimuli (C>P) have a stronger attentional bias towards drug-related cues^7^ and, when trying to quit smoking, are more likely to relapse than individuals with larger neuroaffective responses to pleasant stimuli than to drug-related cues (P>C).^8–10^ However, our previous studies did not directly link the C>P and P>C neuroaffective profiles to cue-induced drug self-administration. Accordingly, this study tested the hypothesis that smokers characterized by the C>P profile are more prone to cue-induced nicotine self-administration than smokers characterized by the P>C profile.

We used the “cued nicotine availability task,” a neurobehavioral laboratory assessment that provides both a neurophysiological measure of the motivational salience attributed to cues signaling nicotine availability and a behavioral measure of cue-induced nicotine self-administration. During the task, event-related potentials (ERPs) are recorded while e-cigarette– naïve smokers look at non-drug-related motivationally relevant images and images of people using an electronic nicotine delivery system (ENDS). The ENDS images signal that an ENDS placed within arm’s reach of the participant is available for use. The cues’ motivational salience is measured using the amplitude of the late positive potential (LPP). The LPP is a component of the ERPs that is higher for stimuli with high motivational salience (e.g., erotic images) than for stimuli with lower motivational salience (e.g., romantic images).^11^ Brain responses to non-drug-related motivationally salient images provide the “motivational context” within which reactivity to drug-related cues can be interpreted.^6^ In previous studies, we showed that multivariate classification algorithms identify two neuroaffective reactivity profiles: in one the LPPs to drug-related cues are higher than those evoked by non-drug-related pleasant images (C>P) and in the other the LPPs to non-drug-related pleasant images are higher than drug-related cues (P>C).^7,9,12^

For this study, we hypothesized that a) we could identify C>P and P>C profiles in smokers by applying cluster analysis to the LPP responses from the cued nicotine delivery task and b) smokers characterized by the C>P profile would use the ENDS to self-administer nicotine significantly more often than those characterized by the P>C profile.

## Methods

### Participants

We enrolled current cigarette smokers who did not use e-cigarettes, were older than 18, and were free from psychiatric disorders. We reimbursed participants with a $50 gift card. The original recruitment plan included 60 smokers. However, owing to the COVID-19 pandemic, our laboratory shut down in March 2020. After one year of inactivity, the protocol associated with this study was closed and only 36 participants were available for the analyses. **Supplementary Table 1** shows the characteristics of the sample.

### Procedures

The University of Texas MD Anderson Cancer Center Institutional Review Board approved all procedures. We asked participants to continue smoking at their regular rate but to refrain from using other drugs (including marijuana) in the 24 hours and caffeine in the 2 hours preceding the visit. At the visit, we explained the procedures and collected informed consent. Then, after collecting self-reports about nicotine dependence, smoking urges, impulsivity, hedonic capacity, and mood, we administered the cued nicotine self-administration task and conducted the electroencephalogram (EEG) assessment. At the end of the session, participants were debriefed, encouraged to quit smoking, and compensated.

## Materials

### Cued nicotine self-administration task

The participants viewed 300 images divided into 6 equivalent testing blocks. The images (selected from the International Affective Picture System^13^ and other collections; see **Supplementary Table 2**) belonged to 1 of 8 categories: erotica (PH, pleasant high salience), romantic (PL, pleasant low salience), food (FD, sweet palatable foods), neutral (NE, ordinary objects and people engaged in mundane activities), unpleasant objects (UO, accidents and pollution), unpleasant low salience (UL, sadness and violence), unpleasant high salience (UH, mutilations), and ENDSs (EC). The images (except EC) were presented for 2 seconds, followed by a 1.5-to 3-second variable inter-trial interval. EC images were presented for 1 second, and then a banner at the top of the screen indicated that it was possible to take 1 puff from the ENDS (Model: THERION-BF-DNA75C, loaded with tobacco-flavored e-liquid with 1.2% nicotine). The ENDS was inside a receptacle within arm’s reach of the participant. A photocell detected when the participant picked up the ENDS and paused the task until the ENDS was placed back in the receptacle. If the participant decided not to use the ENDS, they pressed a button to move to the next trial. Each block included 10 vaping opportunities. To familiarize participants with the ENDS, we ran a practice block of 10 trials with 2 vaping opportunities. We asked participants to take 1 puff at both vaping opportunities.

### EEG data acquisition

During the task, we collected EEG using a 129-channel Geodesic Sensor Net, amplified with a Geodesic EEG System 400 amplifier. The sampling rate was 250 Hz, and all electrodes were referenced to Cz.

### EEG data reduction and LPP amplitude calculation

We reduced the EEG data by following a standard pipeline.^9,14^ It included filtering (0.1-30 Hz), EEG visual inspection and broken channel interpolation (using spherical splines), average reference calculation, and eye movements and blink correction (as implemented in BESA 5.3). Then, data were imported into Brain Vision Analyzer and divided in 1000-ms segments starting 100 ms before picture onset. After baseline correction, channels contaminated by artifacts were identified using pre-defined criteria of relative and absolute voltage amplitude. Segments with more than 10% of contaminated channels were discarded, and average amplitudes for each picture category were computed at each scalp site. For each participant, we calculated the mean LPP amplitude of each category as the mean voltage recorded between 400 and 800 ms after picture onset across 10 central and parietal sensors (EGI HydroCel Geodesic Sensor Net sensors: 7, 31, 37, 54, 55, 79, 80, 87, 106, 129; see **Figure 1** inset).

**Figure 1.**
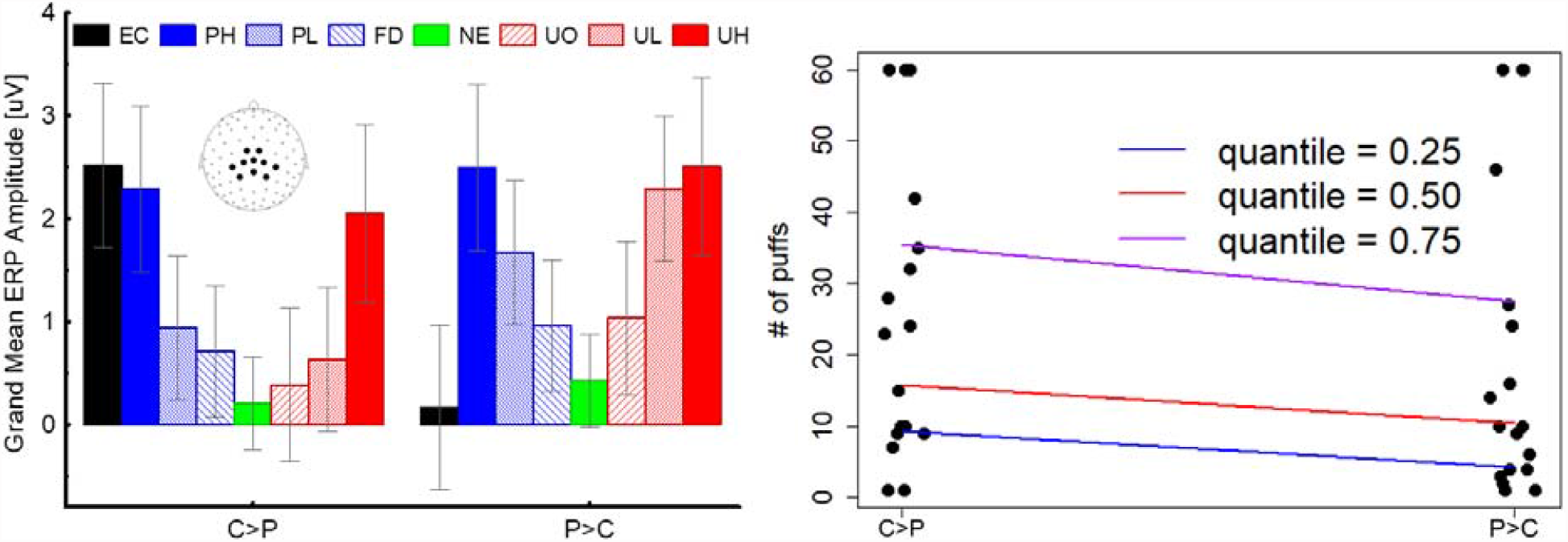
**Left:** The two groups identified by cluster analysis showed the hypothesized characteristics: one group (C>P) had larger late positive potential (LPP) responses to EC than to pleasant images and the other (P>C) had larger LPP responses to pleasant than to EC images. **Right:** Individuals categorized as C>P took significantly more puffs from the electronic nicotine delivery system (ENDS) than individuals categorized as P>C. The between-groups difference in number of puffs was significant (p<.05) at every predicted quantile (.25, .50, and .75). Note: EC=ENDS, PH=pleasant high motivational salience, PL=pleasant low motivational salience, FD=food, NE=neutral, UO=unpleasant objects, UL=unpleasant low motivational salience, UH= unpleasant high motivational salience. The values are calculated averaging the voltage across the 10 sensors shown in the inset.

### Statistical Analyses

#### Event-related potentials

As a manipulation check, we used the LPP responses to the 8 stimulus categories (EC, PH, PL, FD, NE, UO, UL, UH) in a repeated measures ANOVA. We expected to replicate previous findings showing that both pleasant and unpleasant images increase the LPP amplitude as a function of their motivational relevance and that nicotine cues prompt larger LPP responses than neutral stimuli.

#### Participant classification

To identify participants attributing high or low salience to cues, we applied *k*-means cluster analysis to the LPP responses evoked by the 8 image categories. *K*-means cluster analysis is a multivariate classification procedure that, by minimizing within-group variability and maximizing between-group variability, groups individuals according to common features. On the basis of previous findings,^6,7,12^ we hypothesized that the 2 groups will have the following characteristics: 1 group will be characterized by high reactivity to EC relative to pleasant stimuli (C>P) and the other will be characterized by low reactivity to EC relative to pleasant stimuli (P>C). We also expected that, irrespective of reactivity to EC, both groups would show larger LPP responses as a function of motivational relevance for both pleasant and unpleasant stimuli.

#### Cue-induced nicotine self-administration

We tested the relationship between group membership and number of puffs using quantile regression. Quantile regression examines percentiles of an outcome variable distribution. Here, we compared the median (50th), 25th, and 75th percentiles of the number of puffs by group. Since the number of puffs is a count measure, we used quantile regression for counts.^15^ Quantile regression for counts overcomes the issue of non-differentiable sample objective function with discrete dependent variables through artificially imposing smoothness by adding a uniformly distributed noise to the data. A further advantage is that quantile regression is more robust to outliers than least squares regression and does not assume a parametric distribution of the error process. ^16^ For estimation, we used 5,000 bootstrap samples for the quantile regression coefficients, and 95% confidence intervals (CIs) were calculated.

## Results

### Late Positive Potential

**Supplementary Figure 1** shows the grand-averaged ERP waveforms and the mean LPP amplitude for each category. As expected, pleasant, unpleasant, and nicotine-related stimuli prompted larger LPPs than neutral images (Bonferroni-corrected P values <0.005), and the amplitude of the LPP increased as a function of motivational relevance for both pleasant and unpleasant stimuli (polynomial contrast for the quadratic trend including PH, PL, FD, NE, UO, UL, UH; F(1,35)=123.3; p<0.001).

*K*-means cluster analysis assigned 18 subjects to each group. As hypothesized (see **Figure 1**), both groups had larger LPP responses to motivationally salient images than to neutral images (the quadratic trend was significant [p<0.001] in both groups), but 1 group showed the C>P profile (i.e., larger LPP responses to EC than to pleasant images), while the other showed the P>C profile. The 2 groups did not differ according to demographic characteristics, nicotine dependence, mood, or impulsivity (see **Supplementary Table 1**).

### Cue-induced nicotine self-administration

The results of the quantile regression analysis for counts are presented in **Figure 1** and **Table 1**. The unstandardized coefficients do not have an intuitive interpretation because they are derived from a log-linear model. Hence, we estimated marginal effects that have a direct interpretation. Marginal effects are the differences in the number of puffs between the groups derived from each quantile regression model. For all 3 estimated quantiles, the C>P group took a significantly higher number of puffs than the P>C group. Specifically, the differences in the estimated number of puffs were 5 (C>P = 9 vs. P>C = 4), 5 (C>P = 15 vs. P>C =10),, and 8 (C>P = 35 vs. P>C = 27), puffs at the 25^th^, 50^th^, and 75^th^ percentiles, respectively.

**Table 1.** Unstandardized and marginal effects of group membership on the number of puffs.

| Model | Effect | CI | p-value |
| --- | --- | --- | --- |
| **Unstandardized Effects** |  |  |  |
| Quantile 0.25 | -0.80 | (-1.29, -0.31) | 0.001 |
| Quantile 0.50 | -0.41 | (-0.76, -0.07) | 0.018 |
| Quantile 0.75 | -0.26 | (-0.43, -0.09) | 0.002 |
| **Marginal effects of differences between groups** |  |  |  |
| Quantile 0.25 | -5.00 | (-7.61, -2.39) | < 0.001 |
| Quantile 0.50 | -5.08 | (-9.66, -0.49) | 0.029 |
| Quantile 0.75 | -7.99 | (-12.95, -3.03) | 0.002 |

## Discussion

By applying cluster analysis to neuroaffective responses evoked by a wide array of motivationally salient stimuli, we identified smokers who attribute high salience to cues indicating nicotine availability (C>P), and we showed that these individuals are more prone to cue-induced nicotine self-administration than those who do not attribute high salience to such cues (P>C).

Preclinical findings^5^ and our previous results^7,9,14^ indicate that individual differences in the tendency to attribute motivational salience to cues predicting rewards are a transdiagnostic biomarker associated with vulnerability to cue-induced compulsive behaviors. This biomarker could be targeted to match existing treatments to the individuals most likely to benefit from them ^17^ and to develop new tailored treatments for substance use disorders and other behavioral disorders characterized by cue-induced compulsive behaviors.^18^

Notwithstanding their potential significance, these results should still be considered preliminary. A major limitation of this study is its small sample size. We plan to collect a larger sample to ensure the replicability of the effects reported here and to investigate the role that cognitive control has in regulating cue-induced drug seeking.^19^

Previous brain imaging studies investigated the relationship between neurophysiological responses to drug-related cues and craving.^20^ While craving is considered a clinically significant symptom,^21^ it is not consistently associated with relapse.^22^ These inconsistencies indicate that self-reported craving is not a good surrogate measure of cue-induced drug use. By relying on behaviors rather than self-reported craving, the cued nicotine availability task can more directly contribute to identifying the psychophysiological underpinnings of cue-induced drug use. Assessing the extent to which laboratory cue-induced self-administration predicts real-world relapse will contribute to the development and optimization of new targeted clinical interventions for substance use disorders.

## Acknowledgements

This work was supported by the National Institute on Drug Abuse of the National Institutes of Health (R01DA032581 to FV) and by MD Anderson’s Cancer Center Support Grant P30CA016672. The content is solely the responsibility of the authors and does not necessarily represent the official views of the National Institutes of Health. We thank Liz Lee, Kendra Lumbi, and Menton Deweese for help with data collection. The data supporting our findings are available upon request.

## SUPPLEMENTARY MATERIALS

### Self-report questionnaires used in the study

#### Fagerström Test for Cigarette Dependence (FTCD)

The FTCD ^1^ is a 6-item questionnaire that measures nicotine dependence by assessing various components of smoking behavior such as daily intake, difficulty in refraining from smoking, and time to first cigarette. Higher scores indicate higher dependence.

#### Questionnaire of Smoking Urges (QSU-brief)

The QSU-brief ^2^ is a 10-item questionnaire that measures craving to smoke. In addition to a total score, two scores can be extracted: one associated with strong desire and intention to smoke and smoking being perceived as rewarding; the other associated with anticipation of relief from negative affect and an urgent desire to smoke.

#### Positive and Negative Affect Scale (PANAS)

The PANAS ^3^ comprises two 10-item mood scales: Positive Affect (PA) and Negative Affect (NA). Participants rated different feelings and emotions on a scale of 1-5 using “The past few days” as a time reference.

#### Snaith-Hamilton Pleasure Scale (SHAPS)

The SHAPS ^4^ is a 14-item self-report instrument designed to assess hedonic tone that was specifically developed to be unaffected by social class, gender, age, dietary habits, or nationality. Higher scores indicate higher inability to experience pleasure. We scored the answers using a 0-3 Likert scale. Thus the score range is 0-42.

#### Barratt Impulsiveness Scale (BIS)

The BIS ^5^ is a 30-item self-report instrument designed to assess impulsiveness. The participant rates on a 4-point scale the frequency of behaviors and preferences associated with impulsiveness. Higher scores indicate higher impulsiveness on 3 sub-traits: “Attentional Impulsiveness”, “Motor Impulsiveness”, “Nonplanning Impulsiveness”.

### Participants

**Supplementary Table 1.** Demographic characteristics

|  | All (N=36) | C>P (N=18) | P>C (N=18) |
| --- | --- | --- | --- |
| Mean age (SD) | 46 (11) | 42 (12) | 49 (10) |
| Females | 42% | 39% | 44% |
| Race |  |  |  |
| White | 17% | 22% | 11% |
| Black | 83% | 78% | 89% |
| **Questionnaire scores** |  |  |  |
| FTCD | 4.7 (4.7) | 5.2 (1.6) | 4.2 (2.1) |
| QSU | 47.1 (29.4) | 53.2 (31.9) | 50.1 (30.4) |
| BIS<sub>attention</sub> | 14.9 (3.2) | 14.4 (3.4) | 15.4 (3) |
| BIS<sub>motor</sub> | 22.2 (4.7) | 21.4 (4.6) | 23.1 (4.8) |
| BIS<sub>nonplanning</sub> | 24.5 (4.3) | 23.9 (3.3) | 25.1 (5.1) |
| PANAS<sub>negative</sub> | 17.6 (5.4) | 17.7 (5.2) | 17.6 (5.7) |
| PANAS<sub>positive</sub> | 34 (8.7) | 33.9 (9.2) | 34 (8.5) |
| SHAPS | 9.9 (5.5) | 10.6 (5.4) | 9.2 (5.6) |

NOTE: FTCD = Fagerström Test for Cigarette Dependence, QSU=Questionnaire for Smoking Urges, BIS = Barratt Impulsiveness Scale, PANAS=Positive and Negative Affect Scale, SHAPS=Snaith-Hamilton Pleasure Scale.

### Images used in the study

**Supplementary Table 2.** reports the IAPS numbers of the pictures that we used in the experiment. When the IAPS did not have enough pictures for a specific category, we downloaded from the internet images with similar characteristics. All images depicting electronic delivery systems were downloaded from the internet.

| Category | IAPS numbers |
| --- | --- |
| PH | 4604, 4611, 4650, 4653, 4658, 4659, 4660, 4668, 4669, 4677, 4680, 4687, 4690, 4691, 4693, 4694, 4695, 4696, 4698, 4800, 4647-2, 4666-2, plus 8 images not in the IAPS |
| PL | 2208, 2501, 2550, 4597, 4599, 4600, 4610, 4612, 4616, 4619, 4624, 4625, 4640, 4641, 4643, 4700, plus 14 images not in the IAPS |
| PS | 7330, 7390, 7405, 7410, 7430, 7470, plus 24 images not in the IAPS |
| NE | 2037, 2039, 2102, 2107, 2190, 2191, 2210, 2273, 2305, 2359, 2374, 2377, 2383, 2393, 2396, 2397, 2411, 2435, 2441, 2500, 2511, 2512, 2575, 2594, 2595, 2620-2, 2630, 2635, 7550, 9070, 5390, 7000, 7001, 7002, 7006, 7009, 7010, 7011, 7012, 7018, 7020, 7021, 7026, 7030, 7034, 7040, 7041, 7050, 7052, 7053, 7054, 7055, 7056, 7059, 7061, 7062, 7081, 7090, 7150, 7233. |
| UO | 6020, 7079, 7521, 9010, 9090, 9110, 9290, 9291, 9295, 9300, 9301, 9320, 9322, 9373, 9560, 9600, 9621, 9902, 9911, 9912, 9941, plus 9 images not in the IAPS |
| UL | 2703, 2811, 3500, 3530, 6210, 6211, 6230, 6231, 6242, 6260, 6312, 6313, 6315, 6350, 6360, 6510, 6540, 6550, 6560, 6571, 6832, 9414, 9520, 9530, 9429-2, plus 5 images not in the IAPS |
| UH | 3000, 3030, 3051, 3053, 3060, 3064, 3068, 3069, 3071, 3080, 3100, 3103, 3110, 3120, 3130, 3140, 3150, 3170, 3180, 3181, 3211, 3213, 3225, 3261, 3400, 6021, 9253, 9265, plus 2 images not in the IAPS |

### RESULTS

**Supplementary Figure 1.**
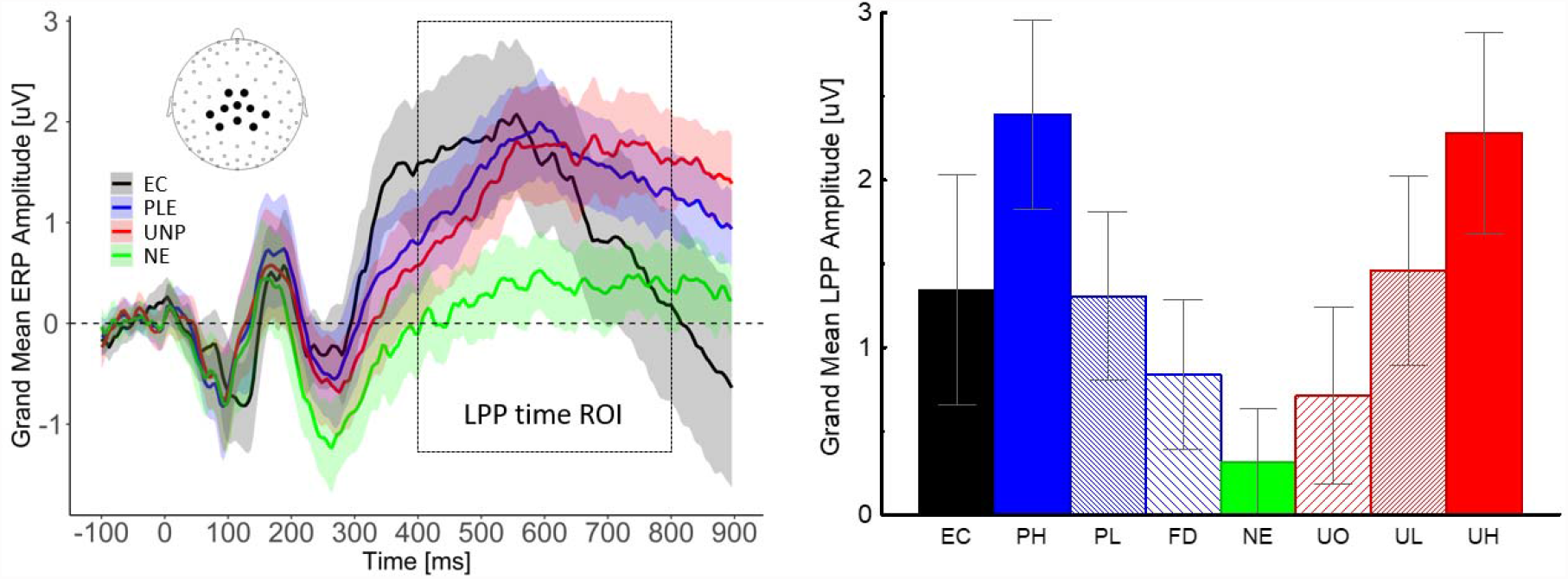
**Left**: As expected, motivationally salient images (including EC) prompted larger LPPs than neutral images. **Right:** The LPP amplitude increased as a function of motivational salience for both pleasant and unpleasant contents. **NOTE**: LPP= late positive potential, ROI=region of interest, EC= E-cigarettes, PLE= pleasant (PH, PL, FD averaged), UNP=unpleasant (UH, UL, UO averaged), NE= neutral, PH=pleasant high motivational salience (erotica), PL=pleasant low motivational salience (romantic), FD=food (appetizing sweet food), UO=unpleasant objects (accidents, pollution), UL=unpleasant low motivational salience (violence), UH= unpleasant high motivational salience. The values are calculated averaging the voltage across the 10 sensors shown in the inset on the left panel.

